# Single-cell identity definition using random forests and recursive feature elimination

**DOI:** 10.1101/2020.08.03.233650

**Authors:** Madeline Park, Sevahn Vorperian, Sheng Wang, Angela Oliveira Pisco

**Affiliations:** Chan Zuckerberg Biohub, San Francisco, California, USA; Department of Chemical Engineering, Stanford University, Stanford, California, USA; Department of Bioengineering, Stanford University, Stanford, California, USA; Department of Genetics, Stanford University, Stanford, California, USA

## Abstract

Single-cell RNA sequencing (scRNA-seq) enables the detailed examination of a cell’s underlying regulatory networks and the molecular factors contributing to its identity. We developed scRFE with the goal of generating interpretable gene lists that can accurately distinguish observations (single-cells) by their features (genes) given a metadata category of the dataset. scRFE is an algorithm that combines the classical random forest classifier with recursive feature elimination and cross validation to find the features necessary and sufficient to classify cells in a single-cell RNA-seq dataset by ranking feature importance. It is implemented as a Python package compatible with Scanpy, enabling its seamless integration into any single-cell data analysis workflow that aims at identifying minimal transcriptional programs relevant to describing metadata features of the dataset. We applied scRFE to the Tabula Muris Senis and reproduced established aging patterns and transcription factor reprogramming protocols, highlighting the biological value of scRFE’s learned features.

**Author summary:** scRFE is a Python package that combines a random forest classifier with recursive feature elimination and cross validation to find the features necessary and sufficient to classify cells in a single-cell RNA-seq dataset by ranking feature importance. scRFE was designed to enable straightforward integration as part of any single-cell data analysis workflow that aims at identifying minimal transcriptional programs relevant to describing metadata features of the dataset.

## Introduction

Single-cell RNA sequencing (scRNA-seq) facilitates newfound insights into the long withstanding challenge of relating genotype to phenotype by resolving the tissue composition of an organism at single-cell resolution [1]. Moreover, scRNA-seq enables the delineation of the constituents contributing to the homeostasis of an organism by understanding the widely varying functional heterogeneity of every cell [2].

Tremendous efforts have been put towards developing methods for cell reprogramming both computationally and experimentally. Cell types are historically characterized by morphology and functional assays, but one of the main advantages of single-cell transcriptomics is their ability to define cell identities and their underlying regulatory networks [30]. As an example, transcription factors (TFs) mediate cell fate conversions [3, 14], which could address the need for specific cell types in regenerative medicine and research [5,8]. A panoply of biological experiments has been performed to further validate and understand the role that transcription factors play in determining cell subpopulations, demonstrating that it is possible to transform the fate of a cell with a unique group of TFs [6, 7]. Furthermore, several computational methods exist for finding those particular sets of TFs [8]. However, these models often rely on an exceptionally well-characterized target cell type population and a ‘background population’ [8], hindering generalizability. These approaches are limited by noisy data and high sensitivity to technical differences in experimentation methods, yielding a large list of TFs to be considered and hampering their usability by bench scientists [8].

Here we present scRFE (single-cell identity definition using random forests and recursive feature elimination, pronounced ‘surf’), a method that combines the classical random forest algorithm with recursive feature elimination and cross validation to find the features necessary and sufficient to classify single-cell RNA-seq data by ranking feature importance. A random forest is an ensemble of uncorrelated decision trees, where for a given split in a given tree, a random subset of the feature space is considered [11]. Random forests are ideal for single-cell data given their effective performance on sparse, high dimensional data with collinear features and straightforward understandability [11].

The goal of scRFE is to produce interpretable gene lists for all cell types in a given dataset that can be used to design experiments or to further aid with biological data mining. The out-of-the-box compatibility of scRFE with Scanpy [9] enables its seamless integration into any single-cell data analysis workflow that aims at identifying minimal sets of genes relevant to describing metadata features of the dataset. Thus, scRFE has potential applications not only in finding transcription factors suitable for reprogramming analysis, but more generally in identifying gene sets and their importances for a given subpopulation of cells.

## Results

### scRFE Overview

scRFE (Fig. 1) takes as input an AnnData object and a class of interest which corresponds to a metadata column of the dataset. The algorithm iterates over the set of labels (specific values) in the class of interest (category to split observations by). scRFE utilizes a random forest with recursive feature elimination and cross validation to identify each feature’s importance for classifying the input observations. Downsampling is performed separately for each label in the class of interest, ensuring that the groups are balanced at each iteration (implementation described in Methods). In this work, each observation is a single-cell, each feature is a gene, and the data are the expression levels of the genes in the entire feature space. In order to learn the features to discriminate a given cell type from the others in the dataset, scRFE was built as a one versus all classifier. Recursive feature elimination was used to avoid high bias in the learned forest and to address multicollinearity [26]. In this way, scRFE learns the features for a given label in the class of interest and returns the mean decrease in Gini score for the top learned features. This is a numerical measure of the feature’s importance in the entire model by quantifying node purity [16]. Recursive feature elimination removes the bottom 20% of features with the lowest mean decrease in Gini scores at each iteration. We incorporated k-fold cross validation (default = 5), to minimize variance and reduce the risk of overfitting [12]. scRFE returns the Gini importances [16] ranking the top features for each label in the class of interest, which are the selected features necessary and sufficient to describe the metadata feature of the dataset due to the implementation of recursive feature elimination and cross validation.

**Fig. 1:**
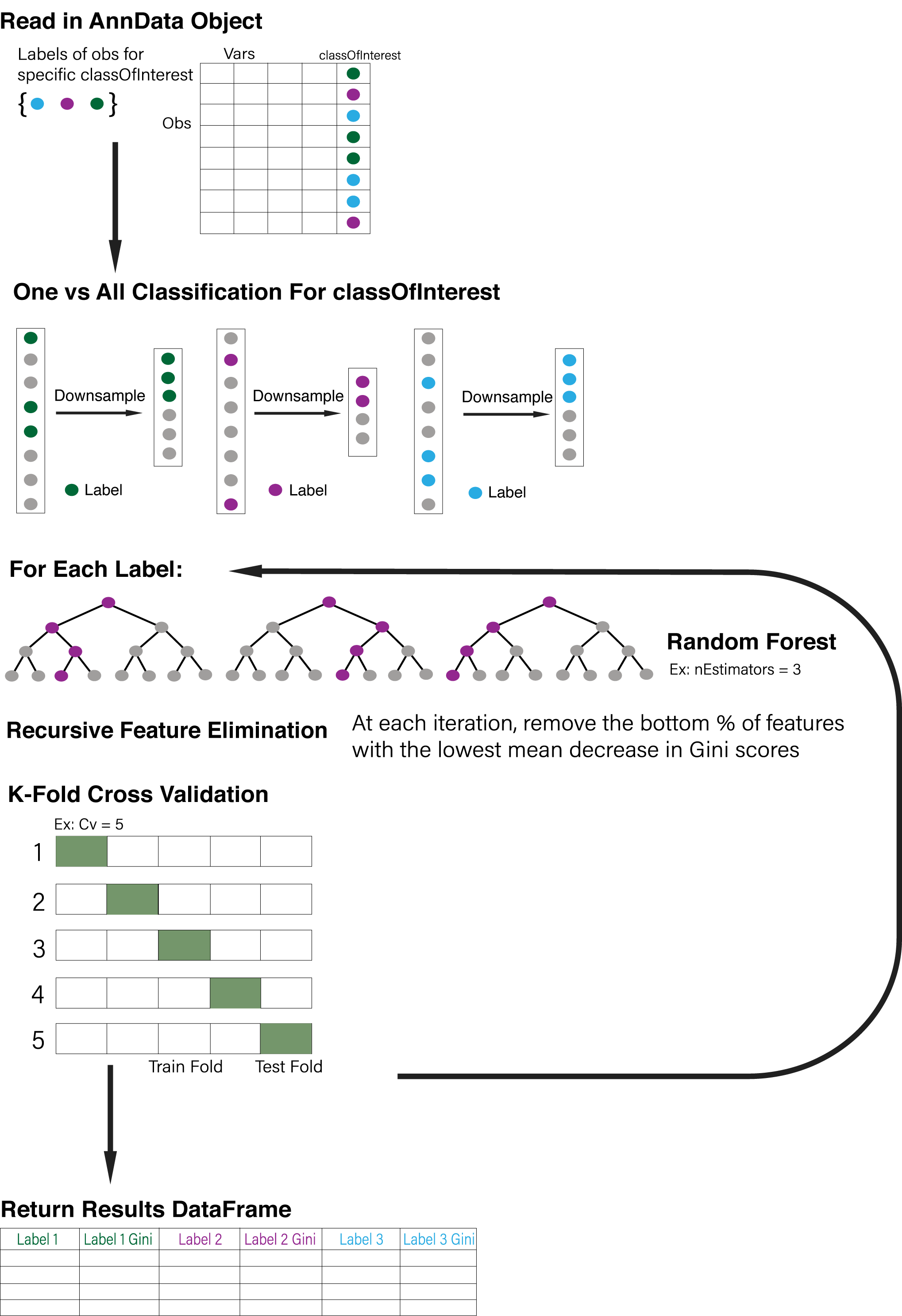
scRFE overview. Given an input AnnData object and specified value for ‘classOfInterest,’ the metadata category of the dataset to split observations by, scRFE performs one vs all classification. scRFE downsamples to the number of observations in smallest category to provide an equal sized comparison for every label within the classOfInterest. For each label, scRFE learns a random forest with recursive feature elimination, where the bottom percent (default = 20%) of features were removed at each iteration. scRFE leverages k-fold cross validation (default k = 5). scRFE outputs a pandas dataframe with columns corresponding to each label within the classOfInterest, storing the selected features and their respective mean decrease in Gini importances.

### scRFE identifies concordance between FACS and droplet techniques in Tabula Muris Senis

We have applied scRFE to the Tabula Muris Senis (TMS) dataset [13]. TMS is a comprehensive atlas of cells from 23 tissues over the murine lifespan. The dataset’s deep characterization of the cell types comprising these tissues over time allowed us to explore TF reprogramming protocols for a diverse range of cell ontologies [14] and aging signatures [15] on the scale of hundreds of thousands of cells.

Comparing the top-performing genes for both the 10X/droplet and Smart-Seq2/FACS datasets from TMS indicates that scRFE’s performance is independent of technical aspects of experimentation (Fig. 2, Supplementary Table 1). Looking at the tissue-specific results (Supplementary Table 2), we saw that scRFE output on the mammary gland-specific FACS and droplet objects with age as the class of interest outputs *≥*58% of shared genes for the 3-month mice (Fig. 2a). scRFE run on the lung-specific FACS and droplet objects with age as the class of interest outputs *≥*61% of shared features for the 18-month old mice (Fig. 2b). This finding remained true when scRFE was run on cell types (Supplementary Table 3), as the FACS and droplet liver-specific objects shared *≥*74% of selected features for hepatocyte cells (Fig. 2c). However, perfect overlap between results is still not achieved due to the inherent experimental differences between the FACS and droplet techniques, with the FACS method generating full-length cDNAs with a greater ability to detect rare cells, compared to the droplet method, an end-to-end solution with high cell capture efficiency [20]. A future interesting approach would be combining the independent datasets to create a more robust characterization of the class of interest.

**Table 1.** Examples of scRFE identifying biologically consistent transcription programs.

**scRFE Table 1**
| Cell Type | Transcription Factors<br>*Bolded are ones scRFE found<br>in top 25 selected features | Reference |
| --- | --- | --- |
| Pericyte Cells | Lmcd1, **Foxf2**, **Tbx2**,<br>**Nr1h3**, Nr2f1 | Okawa, S., et al.<br>Nucleic Acids Res 2019. |
| NK Cells | **Tbx21**, **Runx3**, **Gata3**,<br>Ikzf3, Tcf7 | Okawa, S., et al.<br>Nucleic Acids Res 2019. |
| Oligodendrocytes | **Sox10**, **Olig2**, **Zfp536** | Yang, N., et al.<br>Nat Biotechnol 2013. |
| Skeletal Muscle<br>Cells | **Myod1** | Davis, R.L., et al.<br>Cell 1987. |
| Neuronal Stem<br>Cells | **Sox2** | Ring, K.L., et al.<br>Cell Stem Cell 2012. |

**Table 2.**
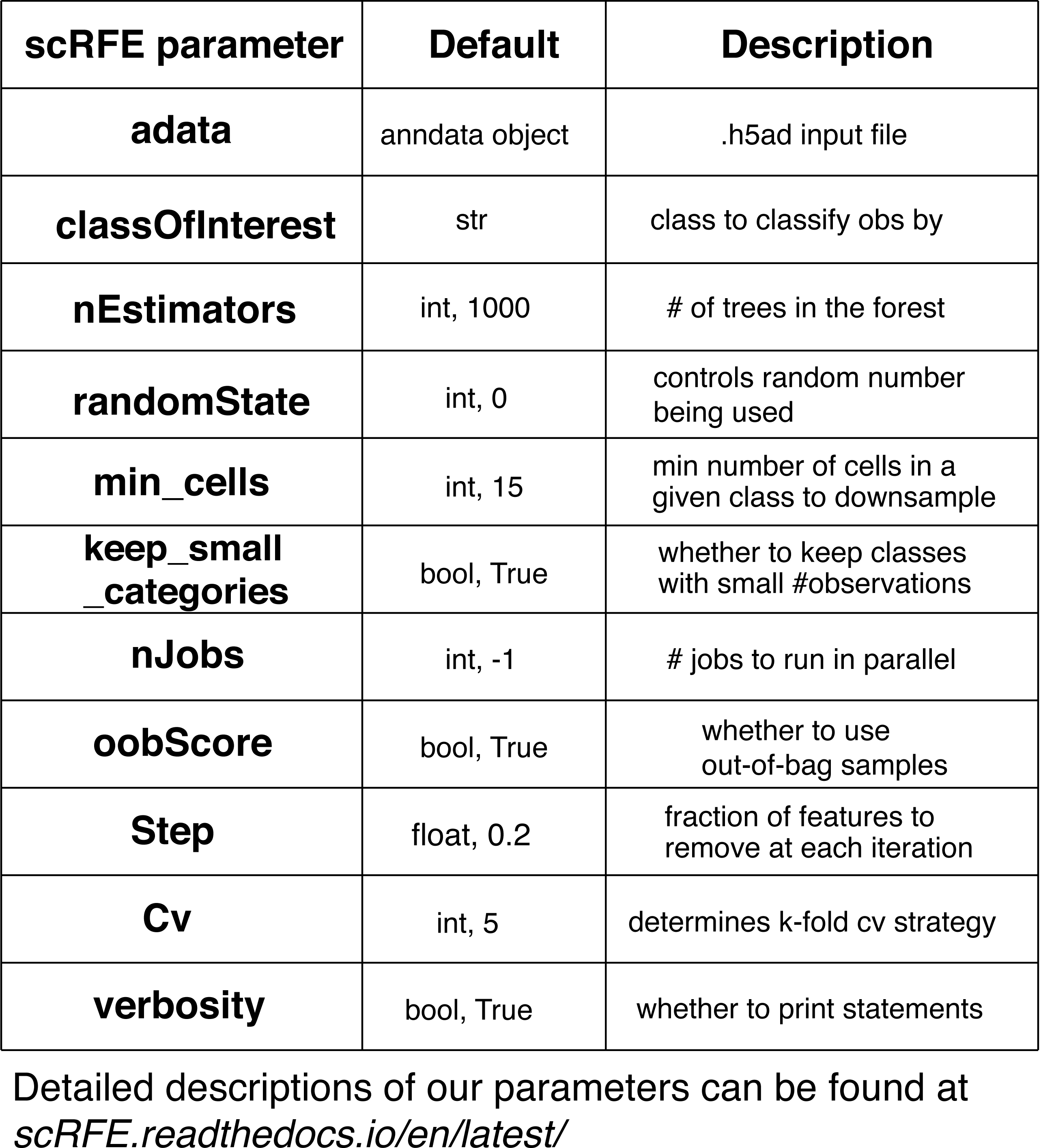
scRFE parameters default values and descriptions.

**Fig. 2:**
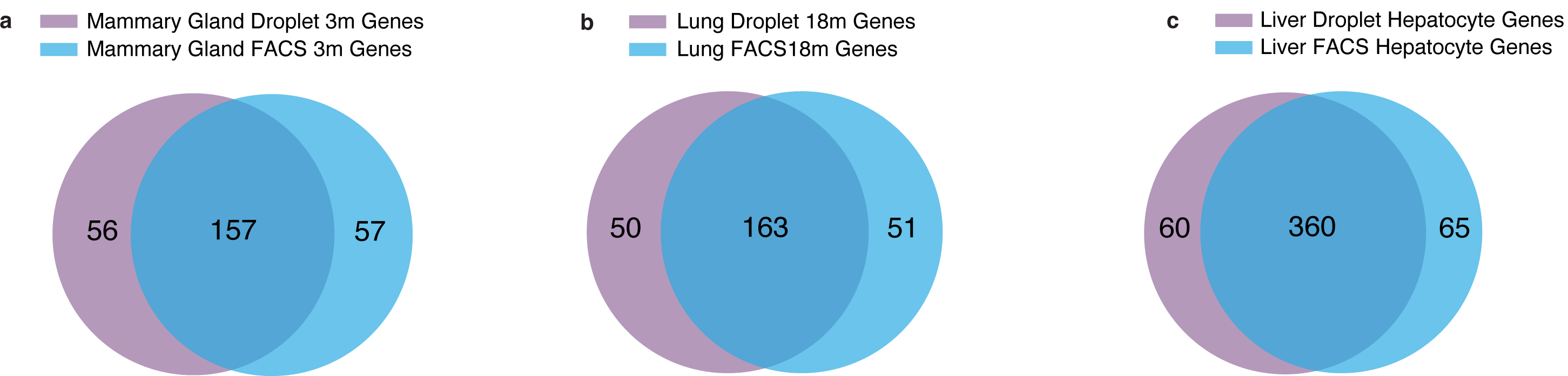
scRFE finds consistency between FACS and droplet methods. Venn diagrams visualizing the intersection of scRFE results with default parameters, run on tissue-specific FACS versus droplet objects. a) Mammary gland with classOfInterest = ‘age.’ Here, we selected the results for 3-month. Majority of the selected genes for both datasets were shared. The two results overlapped *≥*58% of the time with 157 shared genes, 56 droplet specific genes, and 57 FACS specific genes. b) Lung with classOfInterest = ‘age.’ Here, we selected the results for 18-month. Majority of the selected genes for both datasets were shared. The two results overlapped *≥*61% of the time with 163 shared genes, 50 droplet specific genes, and 51 FACS specific genes. c) Liver with classOfInterest = ‘cell ontology class.’ Here, we selected the results for hepatocytes. Majority of the selected genes for both datasets were shared. The two results overlapped *≥*74% of the time with 360 shared genes, 60 droplet specific genes, and 65 FACS specific genes.

### scRFE learns biologically consistent transcriptional programs

Specific sets of transcription factors determine a cell’s fate [3] and our method finds concordance with known marker genes for different cell ontologies coming from a variety of tissues. For example, Lmcd1, Foxf2, Tbx2, Nr1h3, and Nr2f1 are linked to pericyte cells [7]. scRFE found Foxf2, Tbx2,and Nr1h3 in the top four selected features for pericyte cells when run on the lung droplet object, splitting observations by cell type. For NK cells from the same output, scRFE found Tbx21, Runx3, and Gata3 in the top 22 selected features, three of the five TFs linked to NK cells [7]. Additionally, Sox10, Olig2, and Zfp536 are reprogramming factors for oligodendrocytes [17] and scRFE found all three TFs in the top eleven selected features for oligodendrocytes when run on the brain non-myeloid FACS dataset, splitting observations by cell type. Not only was scRFE able to recapitulate sets of genes for identifying TFs (Table 1), but the numerical feature rankings also followed existing literature [7, 17–19]. The features with high mean decrease in Gini scores were often already experimentally validated and/or computationally derived as shown by the colored genes in Fig. 3 [3, 17–19].

**Fig. 3:**
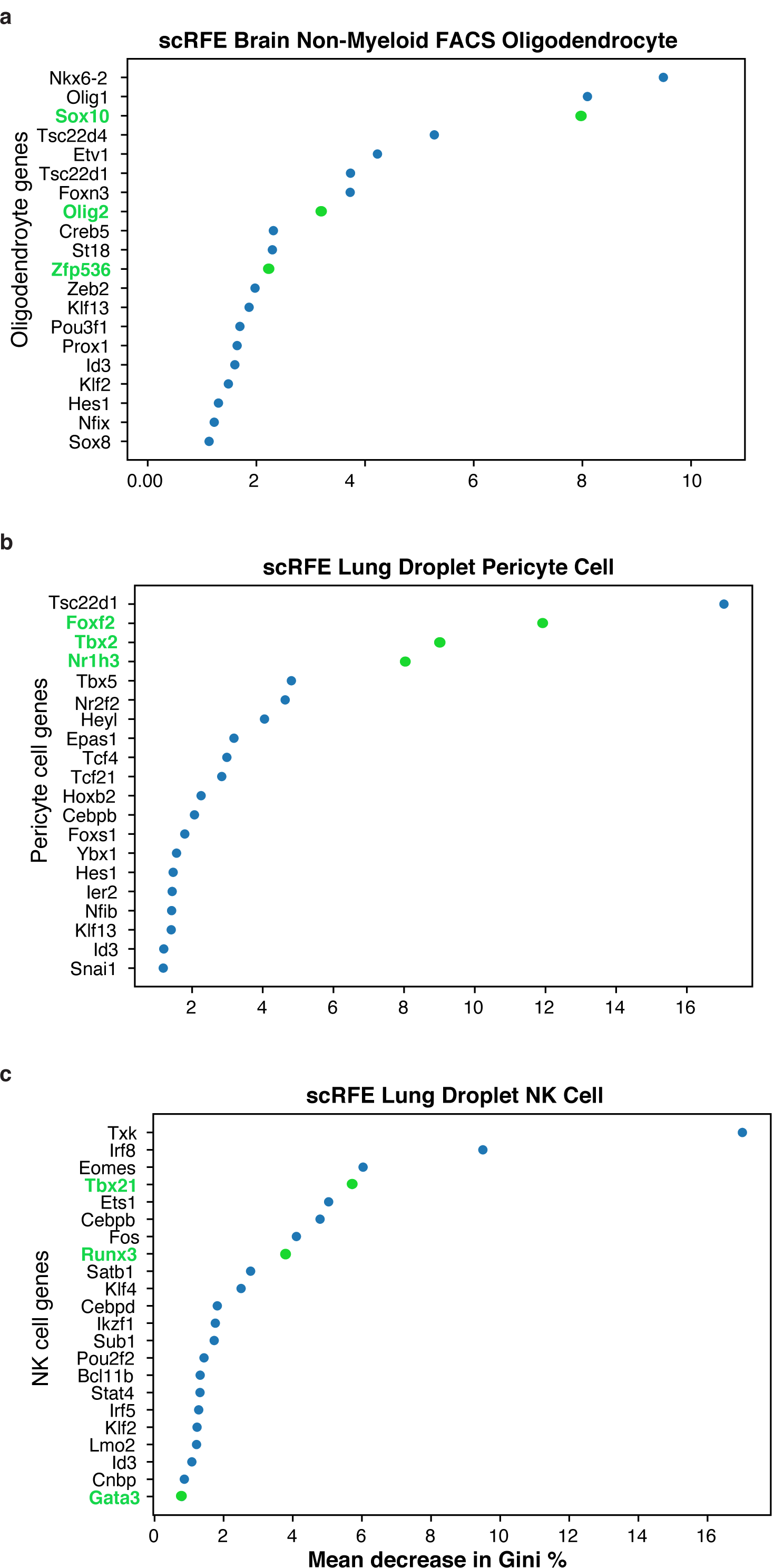
scRFE learns biologically consistent transcriptional programs. a) Sox10, Olig2, and Zfp536 are necessary to reprogram cells into oligodendrocytes [17]. scRFE found all three transcription factors in the top eleven selected features as shown in green. b) ‘Synergistic identity core’ of transcription factors for pericyte cells in the murine liver includes Lmcd1, Foxf2, Tbx2, Nr1h3, and Nr2f1 [7]. scRFE found three out of these five TFs in the top four selected features as shown in green. c) ‘Synergistic identity core’ of transcription factors for NK cells in the murine liver includes Tbx21, Runx3, Gata3, Ikzf3, and Tcf7 [7]. scRFE found three of these in the top 22 selected features as shown in green.

### Top feature importances reveal gene ontology patterns by age

The aging phenotype is associated with notable transcriptional changes [27], and we used scRFE to learn the top ranked genes associated with each age. When considering the unique age-specific genes, we found that Ppp1c is the gene with the highest mean decrease in Gini score that is specific to the 24-month mice and not in the 3- or 18-month lists (FACS dataset). Ppp1cb is associated with aging and mortality [28]. The second-highest ranked gene that is specific to only the 24-month list is Sfpq, a gene linked to diseases such as familial amyotrophic lateral sclerosis (ALS) [4]. Additionally, mutations in Sfpq and the dysregulation of the Sfpq-dependent energy metabolism may be connected to sarcopenia, also known as biological aging [10]. 574 genes (out all the mouse protein coding genes) were shared between the 3-, 18-, and 24-month old mice (Supplementary Table 4).

In order to understand the underlying biology of the scRFE aging results, we performed Gene Set Enrichment Analysis (GSEA) [24, 25] and found that the selected features exclusive for 3-month mice are associated with regular biological processes, such as amino-acid metabolism and cell adhesion (Fig. 4a). When looking at either the 18-month (Fig. 4b) or 24-month (Fig. 4c) selected features, excluding the 3-month selected features, we observed a dramatic switch towards a disease signature. For the 18-month and 24-month lists, the top KEGG pathways are associated with both infections and neurological diseases, reiterating scRFE’s ability to recover biologically relevant signatures in the form of minimal transcriptional sets.

**Fig. 4:**
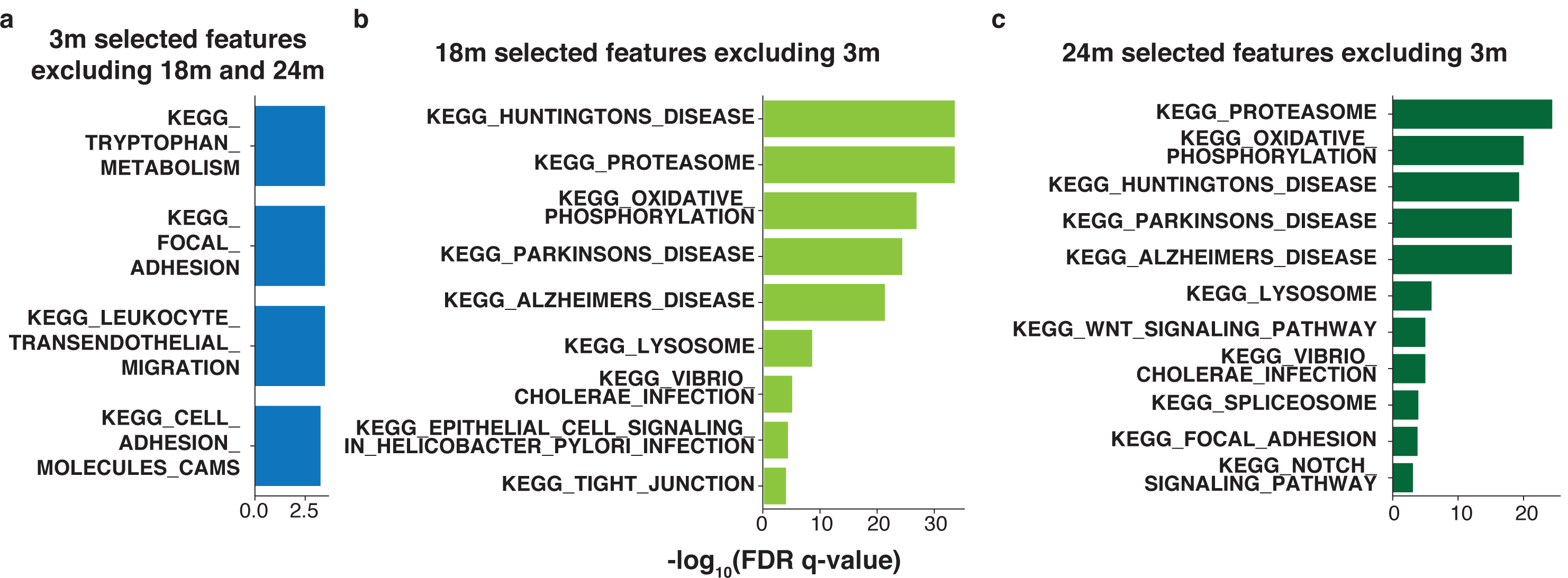
KEGG pathway analysis for scRFE aging results. a) Statistically significant KEGG pathways for the scRFE selected features for 3-month excluding 18-month and 24-month. b) Statistically significant KEGG pathways for the scRFE selected features for 18-month excluding 3-month. c) Statistically significant KEGG pathways for the scRFE selected features for 24-month excluding 3-month.

## Conclusion

scRFE is a robust and reproducible solution to the problem of identifying cell type marker genes and biological transcriptional programs in single-cell RNA sequencing. The code is available on GitHub (https://github.com/czbiohub/scRFE) and it has been included in PyPI (https://pypi.org/project/scRFE/) to facilitate its integration with current scRNA-seq workflows.

When applied to the TMS dataset, scRFE identifies concordance between FACS and droplet techniques, highlighting its potential in detecting biological over technical signals. scRFE also learns transcription factor reprogramming methods which follow biological results. This underscores scRFE’s effectiveness at recognizing reprogramming protocols and potentially finding novel ones.

scRFE allows for the discovery of new associations between genotype and phenotype by finding top marker genes for cell populations. As proof of concept, we demonstrate that scRFE produces results that match commonly known aging patterns, highlighting developmental pathways in young murine models and moving towards canonical aging and disease related pathways in older murine models. Future work may include utilizing scRFE to discover genes and pathways biologically inherent to young versus old organisms, furthermore examining the underlying mechanisms of aging. In sum, scRFE identifies the top genes for a given cell population and validates existing conclusions, underscoring its broad utility as a means of finding novel marker genes.

## Acknowledgments

We thank Jingyi Wei, Jim Karkanias, and Steve R Quake for thoughtful discussions. S.V. is supported by a NSF Graduate Research Fellowship (Grant# DGE 1656518), the Benchmark Stanford Graduate Fellowship, and the Stanford ChEM-H Chemistry Biology Interface (CBI) Training Program. This work was funded by the Chan Zuckerberg Biohub.

## Author’s contributions

M.P. designed and implemented the method, packaged the code for PyPI, performed the computational analysis, and co-wrote the manuscript. S.V. designed and implemented the method and co-wrote the manuscript. S.W. packaged the code for PyPI. A.O.P. conceptualized the study, designed and implemented the method, performed the computational analysis, and wrote the manuscript. All authors read and approved the final manuscript.

## Methods

### Implementation

scRFE is built in Python using methods from the scikit-learn [21] and Scanpy [9] libraries. We chose to use Python over R given its compatibility with Scanpy/AnnData objects. For R users, we recommend using the varSelRF package [22]; example of applications to single-cell dataset in Tabula Muris [14]). scRFE’s direct compatibility with Scanpy makes it easy to integrate as a part of any single-cell data analysis workflow interested in identifying minimal sets of genes relevant to describing metadata features of the dataset.

We define *classOf Interest* as the column of the metadata (adata.obs) that scRFE will split observations by. We call each specific value within the *classOf Interest* a “label.” We refer to the input AnnData object as *adata*. We use the term “mean decrease in Gini score” interchangeably with “Gini importance.”

### Datasets and gene lists

We used Tabula Muris Senis (TMS) for our analysis [13]. TMS comprises the gene expression of over 350,000 cells across 23 organs and tissues from mice aged 1 month to 30 months. By understanding tissue composition at single-cell resolution and the widely varying functional heterogeneity of each cell, a deep exploration of transcription factor reprogramming protocols and aging signatures was possible. TMS was generated using both the FACS and droplet methods which allowed us to explore scRFE’s performance with different experimental techniques.

TMS was downloaded from FigShare (processed official FACS and droplet Ann-Data objects found at https://figshare.com/projects/Tabula Muris Senis/64982). We ran scRFE on the TMS AnnData objects using the processed data. Users may choose to remove cells or genes that do not meet a minimum requirement, or to use raw data prior to running scRFE. The GO mouse transcription factors list was defined as the 1,140 genes annotated by the Gene Ontology term ‘DNA binding transcription factor activity,’ downloaded from the Mouse Genome Informatics database (http://www.informatics.jax.org/mgihome/GO/projects.html).

### scRFE function

scRFE receives *adata*, an AnnData object (.h5ad), as input. The data are used as is in the .*X* attribute. scRFE also receives as input *classOf Interest*, a column of the .*obs* attribute of *adata* that is the category to split observations by.

scRFE begins by removing observations that had a missing (*NaN*) annotation for the *classOf Interest* from the inputted AnnData object. Next, scRFE splits the data for one versus all classification. For every unique label in the specified *classOf Interest*, scRFE iterates through each observation and distinguishes between observations that match the given label of interest in the class, versus observations with other labels. scRFE downsamples to the smallest category, meaning it randomly selected the same number of observations that the *classOf Interest* had from the rest of the observations, allowing for a balanced one versus all comparison.

scRFE sets the *X* value of the random forest to *adata*.*X*, the observations by variables matrix. The *y* value of the random forest are the labels created from the previous step. Then, scRFE uses sklearn to create a random forest (*clf*) and recursive feature eliminator (*selector*). The random forest splits observations (cells) based on what features (genes) are expressed until all observations are classified. Feature importances are calculated using the mean decrease in Gini score, a numerical measurement of node impurity [16].

At each iteration, the selector removes the bottom percent (default = 20%) of features with the lowest mean decrease in Gini importances to avoid high bias and to address multicollinearity [26]. The selector also incorporates k-fold cross validation (default = 5) to minimize variance and reduce the risk of overfitting [12]. Thus, the data are split randomly into 5 folds, where at each iteration, 4 folds are trained. This combination of recursive feature elimination and cross validation allows for not only a robust model to be learned, but an elegant and concise model with only the necessary and sufficient features to describe the data selected. scRFE has several tunable parameters (Table 2) and they should be adjusted given a particular dataset. For the most part, we used scRFE’s default parameters, unless stated otherwise.

### scRFE output

Upon running, scRFE returns two values. The first is a Pandas dataframe with the top ranked features in descending order and their corresponding mean decrease in Gini scores for each label within the *classOf Interest*. This can be saved as a csv or used for downstream analysis as-is. The second value scRFE returns is a dictionary containing the score of the underlying estimator for each label within the *classOf Interest* including only the selected features.

### Running scRFE to generate TF and aging results

We ran scRFE on the TMS [13] processed official FACS and droplet AnnData objects on a variety of combinations of tissues, cell types, and ages. We used scRFE’s default parameters and sweeped the number of trees (scRFE argument nEstimators) to assess model performance. We observed how the results changed with different values of nEstimators to determine 1,000 as our default parameter (Fig. 5).

**Fig. 5:**
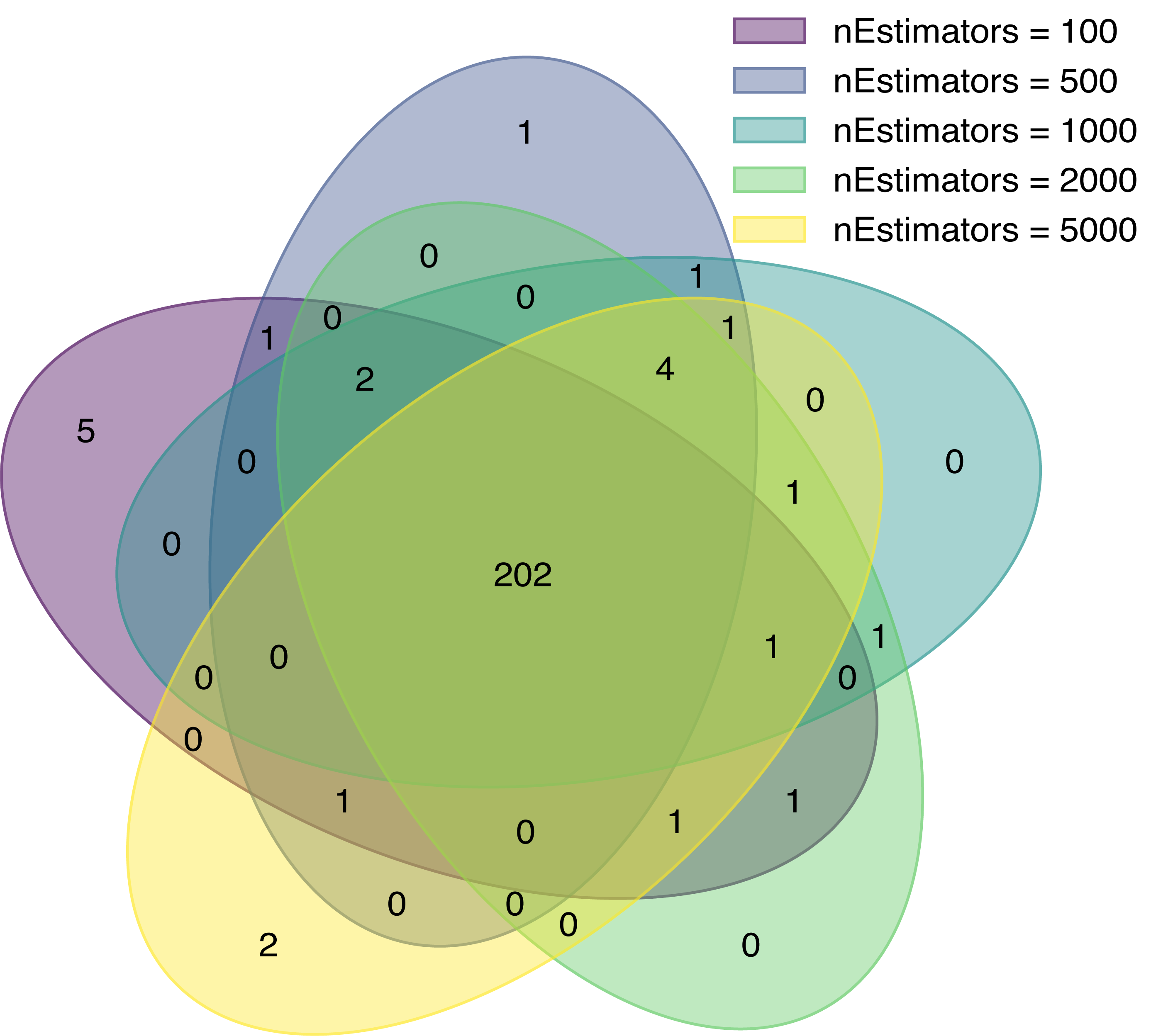
scRFE run on global FACS object with classOfInterest = ‘age’ and different values for nEstimators. 18-month results.

An example of calling scRFE (here, learning top transcription factors per cell type) is presented below; a detailed overview is in the GitHub repository.

~~~
import scanpy as sc
import pandas as pd
from scRFE.scRFE import scRFE
# read in adata and transcription factor list
adata = read_h5ad(“data.h5ad”)
tfs = pd.read_csv(“geneList.csv”)[“Symbol”]
# subset feature space
adataTF = adata[:, tfs]
#call scRFE to split observations by cell type (class of interest)
topFeaturesDF, score = scRFE(adata = adataTF, classOfInterest = “cell_ontology_class”)
~~~

A detailed list of the experiments we discussed in this manuscript can be found in the supplementary tables.

### gProfiler and GSEA analysis

We ran scRFE on the global FACS dataset with age as the *classOf Interest*. For the ages 3-, 18-, and 24-months, we took the lists of features selected and inputted them to gProfiler [23] (Supplementary Table 5). This returned a dataframe highlighting significant pathways (sorted by p-value) for different gene ontologies based on the inputted gene lists (Supplementary Table 6). The intersection of the lists of features selected was analyzed using Gene Set Enrichment Analysis (GSEA MGSig Database) [24, 25] and the results were summarized in Fig.4.

### Code availability

The scRFE code is at https://github.com/czbiohub/scRFE (GitHub). The documentation/user manual is at https://scRFE.readthedocs.io/en/latest/. Pip installation is available at https://pypi.org/project/scRFE/ (PyPI).

## Supplementary Tables

Note: scRFE default parameters were used unless stated otherwise.

**Supplementary Table 1**

scRFE results for FACS and droplet tissue and age-specific objects subsetted for TFs as features with classOfInterest = ‘cell ontology class.’ (cell type).

**Supplementary Table 2**

scRFE results for FACS and droplet tissue-specific objects subsetted for TFs as features with classOfInterest = ‘age.’ nEstimators = 5000.

**Supplementary Table 3**

scRFE results for FACS and droplet tissue-specific objects subsetted for TFs as features with classOfInterest = ‘cell ontology class’ (cell type).

**Supplementary Table 4**

scRFE results for FACS and droplet global objects subsetted for TFs as features with classOfInterest = ‘age.’ nEstimators = 50, 100, 500, 1000, 2000, and 5000.

**Supplementary Table 5**

gProfiler Results for Supplementary Table 2 lists.

**Supplementary Table 6**

scRFE results for FACS global object with classOfInterest = ‘age.’

